# Age-Related Dynamics of Methanogenic Archaea in the Human Gut Microbiome: Implications for Longevity and Health

**DOI:** 10.1101/2024.02.09.579604

**Authors:** Rokhsareh Mohammadzadeh, Alexander Mahnert, Tejus Shinde, Christina Kumpitsch, Viktoria Weinberger, Helena Schmidt, Christine Moissl-Eichinger

**Affiliations:** Diagnostic and Research Institute of Hygiene, Microbiology and Environmental Medicine, Medical University of Graz, Neue Stiftingtalstraße 6, 8010 Graz, Austria; Division of Molecular Biology and Biochemistry, Medical University of Graz, Austria; BioTechMed, 8010 Graz, Austria

## Abstract

The reciprocal relationship between aging and alterations in the gut microbiota is a subject of ongoing research. While the role of bacteria in the gut microbiome is well-documented, specific changes in the composition of methanogens during extreme aging and the impact of high methane production in general on health remain unclear. To address these questions, we analyzed metagenomic data from the stool samples of young adults (n=127, Age: 19-59 y), older adults (n=86), and centenarians (n=34, age: 100-109 years).

Our findings reveal a compelling link between age and the prevalence of high methanogen phenotype, while overall archaeal diversity diminishes. Surprisingly, the archaeal composition of methanogens in the microbiome of centenarians appears more akin to that of younger adults, showing an increase in *Methanobrevibacter smithii*, rather than *Ca.* M. intestini. Remarkably, *Ca.* M. intestini emerged as a central player in the network stability of adults, paving the way for *M. smithii* in older adults and centenarians. Notably, centenarians exhibit a highly complex and stable network of these two methanogens with other bacteria. Furthermore, the mutual exclusion between Lachnospiraceae and these methanogens throughout all age groups suggests that these archaeal communities may compensate for the age-related drop in Lachnospiraceae by co-occurring with butyrate-producing Oscillospiraceae.

This study underscores the crucial role of the archaeal microbiome in human physiology and aging. It highlights age-related shifts in methanogen composition, emphasizing the significance of *Ca.* M. intestini and the partnership between methanogens and specific butyrate-producing bacteria for enhanced health and potential longevity.

## Introduction

Aging is a complex process that affects the physiological, metabolic, and immune functions of humans often leading to chronic inflammation and metabolic issues (1). It is uncertain whether the observed changes in the microbiota are a cause or consequence of aging. According to previous studies, elderly and centenarians tend to have distinct gut microbiome profiles as the latter are able to rearrange the microbiota that contribute to host health and physiology. This includes mitigating the depletion of Ruminococcaceae, Lachnospiraceae, and Bacteroidaceae through the promotion of potential health-enhancing subdominant species like *Akkermansia*, *Bifidobacterium*, and *Christensenellaceae (2-5)*. It is however challenging to determine whether these microbial differences contribute to extreme aging or result from a healthier lifestyle (6). Animal studies suggest that age-related microbial imbalances can impact lifespan, with some animals benefiting from supplementation with a microbiome of younger ones (7) and others experiencing intestinal problems and premature mortality due to aging-associated microbiome changes (8). Despite uncertainties regarding whether gut dysbiosis is a cause or consequence of aging and the subsequent inflammatory disorders, maintaining gut microbiota homeostasis is believed to be crucial for healthy aging and potentially supportive of human longevity (9, 10).

Since short chain fatty acids (SCFAs) produced by the microbiota are absorbed into the host bloodstream through the intestinal epithelium, it is plausible that microbiota-derived metabolites could have a substantial impact on human longevity (11). The decline in metabolic health observed in old age may partly result from altered levels of intestinal SCFAs, and in particular, butyrate, leading to the disruption of gut barrier integrity, increased vulnerability to infections, and affecting conditions like insulin sensitivity and energy expenditure (12). The study by Biagi et al., which is one of the few studies on the microbiome of centenarians residing in Western Europe, revealed the changes in gut microbial composition of these subjects, characterized by a decrease in core abundant taxa like *Bacteroides*, *Roseburia*, and *Faecalibacterium* species, along with an increase in rare taxa. Interestingly, they also observed a change in the population of butyrate-producing bacteria among centenarians. This suggests the possibility that, to achieve longevity, a complex and pervasive remodeling, which includes alterations of gut microbiota, should occur, favoring the balance between inflammatory and anti-inflammatory processes (13).

The human microbiome is not solely composed of bacteria; methanogenic archaea also play a significant but often overlooked role (14). However, our understanding of age-related changes in gut-associated methanogens is limited. It is known that *Methanobrevibacter smithii*, as the predominant archaeal species within the human gut, gradually becomes the dominant archaeal colonizer in early life, with reported higher abundances in centenarians (13). Moreover, Methanomassilicoccales have been frequently observed in elderlies as compared to younger adults (prevalence of 40% vs. 10%) (15), and similarly, a significant age-related upward trajectory was observed for *Methanomassiliicoccus luminyensis* and *Ca.* Methanomassiliicoccus intestinalis (16).

Yet, the specific alterations in the composition of methanogens and their interaction between bacterial members of the human gut during extreme aging remains unknown.

It is noteworthy that a considerable proportion of the human population, approximately 20% of Western adults, falls into the category of high methane emitters, which have been characterized by a 1000-fold increase in *M. smithii* abundance in their gut microbiome. While an association between high methane emissions and complexity within the gastrointestinal microbiome of younger populations has been reported (17), the precise distribution of subjects with high methanogen phenotype and their microbiome across various age groups remains an aspect yet to be elucidated.

Furthermore, recently, the existence of two distinct species within *Methanobrevibacter smithii* has been suggested (18). Indeed, a recent study of archaeal metagenome-assembled genomes (MAGs) underscores the pronounced dissimilarities between *M. smithii_A* and *M. smithii* genomes within the Genome Taxonomy Database (GTDB), to the extent that these distinctions fulfill the threshold criteria for species differentiation, as stipulated by the average nucleotide identity (ANI) metric (>95%). Consequently, *M. smithii_A* has been designated as ‘*Candidatus* Methanobrevibacter intestini’ (18). It remains an open question whether the distribution of *Ca.* M. intestini within the human population varies with age and whether there is a contribution of this archaeal species to methane production, analogous to its counterpart.

In the scope of this study, we sought to discern the diversity and distribution patterns of methanogenic archaea across different age groups of adults. Our study also encompasses a comprehensive examination of the prevalence for a high methanogen phenotype, within varying age groups. In addition, we embark on an exploration of the potential implications and associations of the presence of high methanogen phenotype in the context of extreme longevity. This research provides invaluable insights into the intricacies of archaeal dynamics within the human microbiome, its age-related manifestations, and its putative contributions to the promotion of longevity.

## Materials and Methods

### Study population

To include different age groups, our study incorporated three cohorts. As a young adult cohort, we used fecal samples from 91 subjects enrolled in Graz, Austria, which were initially collected for a study by Kumpitsch et al (Cohort A) (17). To encompass the older adults, a total of 94 participants aged 46-86 (68 ± 9.5) years (female: 51.6%) (Cohort B) were recruited at the Medical University of Graz. The study was evaluated and approved according to the Declaration of Helsinki by the local ethics committee of the Medical University of Graz with the approval number of 26-573 ex 13/14. Before participation, all participants signed informed consent; and finally, to include centenarians (Cohort C) in our study, we chose the metagenomes available in the Sequence Read Archive (SRA) repository under BioProject number PRJNA553191 from the study of Rampelli et al (19), due to the close proximity of the subjects (Emilia Romagna region, Italy) to those enrolled for the first two cohorts.

### Sample collection and DNA extraction

Following collection, stool samples were promptly placed in a stool collection tube and immediately placed on ice. Subsequently, the samples were divided into separate Eppendorf tubes, suspended in a solution of approximately 0.1 gram of fecal material and 0.9% (w/v) DNA-free phosphate-buffered saline, and stored at -20°C for subsequent analyses.

Genomic DNA extraction was carried out on 250 μl of fecal samples using the DNeasy PowerSoil Kit (QIAGEN, USA) according to the manufacturer’s protocol with a slight modification as previously described (Kumpitsch et al.). DNA was eluted in 80 μl elution buffer and the concentration was measured using the Qubit dsDNA HS Assay Kit (Thermo Fisher Scientific, USA).

### Metagenomic sequencing

Extracted DNA from fecal samples, were sent for sequencing to Macrogen (Seoul, South Korea). Library was generated via Nextera XT DNA Library construction kit and sequenced on NovaSeq 6000 Illumina platform. Raw reads were obtained in fastq format with a mean read count of 27,623,362 for each sample.

### Taxonomic classification

Reads were quality checked and human reads were removed as previously described (19). In summary, quality of all reads was assessed with fastqc (v0.11.8) (20), and subsequently filtered with trimmomatic (v0.38) (21) (a minimal length of 50 bp and a Phred quality score of 20 in a sliding window of 5 bp was applied). To filter out the human-mapped reads after quality filtering against the human chromosome GRCh38, bowtie2 (v2.3.5) and samtools (v1.9) were employed (22). Subsequently, bedtools (v2.29.0) was used to extract the fastq files from the bam files (23). We then used Kraken v.2.1.2 (24) to profile these final quality filtered reads with the Unified Human Gastrointestinal Genome (UHGG v.2.0.1) database of genomes, which consists of more than 289.000 archaeal and bacterial genomes. In order to increase the specificity and to compensate for the chance of returning the incorrect lowest common ancestor (LCA) of all genomes, a confidence threshold of 0.3 was chosen for Kraken v.2.1.2. To determine the relative abundance of bacterial and archaeal species, the Kraken2 output was subjected to analysis using Bracken v.2.7. (25) with default settings. The report files were then merged to obtain an abundance table of microbial species which was used for further analysis.

### Removal of the batch effect

Differences in experimental designs and sequencing protocols have the potential to impact the distribution of microbiome data (26). Due to the inclusion of diverse age demographics within three distinct cohorts A, B, and C, and our aim to combine these cohorts for the comprehensive exploration of the microbiome in various age groups, we opted to mitigate the influence of unwanted batch variations arising from distinct study designs and sequencing protocols employed. For this purpose, to generate a batch-corrected table of microbiome read counts for further analyses, we utilized the ConQuR tool (27). This tool effectively addresses read distribution through quantile regression and handles the presence or absence of microbes using conditional quantile regression. We used Cohort B as the reference batch since it removed batch effects the most (the least PERMANOVA R2) and used it across all taxa to keep the overall composition of microbiome and used the presence of high methanogen phenotype (defined based on breath methane test), age classification, and sex as variables.

### Statistical analyses and visualization

Relative and differential abundance of Archaea and bacteria were plotted in R (R-Core-Team, 2022) using the ggplot2 package (v3.3.3). Differentially abundant taxa were defined by q2-ALDex2 (28, 29) in QIIME2 (30). To display those taxa in boxplots in R (packages: ggplot2 (31), dplyr (32), reshape (33)) the data of relative abundance were first CLR (centered log-ratio) transformed in R (34). For statistical analyses, IBM SPSS Amos v26 was used. The normal distribution of parameters were checked using the Shapiro-Wilk test for the selection of the suitable statistical test. Throughout the manuscript, uncorrected significance values are reported as *p*-values and Benjamini-Hochberg corrected *p*-values are termed as *q*-values.

### Co-occurrence analyses

The sparse nature of metagenomic data, which can be attributed to various factors, including sample variations and sequencing depth, presents a challenge when inferring co-occurrence patterns. These variables can introduce challenges in statistical analysis, potentially leading to false-positive results and misleading correlations. To address this issue, we applied a prevalence filter that excluded microbial species present in less than 20% of samples in each age group. We chose this prevalence threshold based on the number of reads for *Ca*. M. intestini to avoid losing this particular species of interest. This approach alleviated the impact of matrix sparsity on our results.

To infer species-level associations of *M. smithii* and *Ca.* M. intestini with other bacteria within the abundance matrix of each age group separately, we employed SparCC (35) within the SCNIC tool (Sparse Co-occurrence Network Investigation for Compositional data) (36). This method for inferring microbial associations incorporates the compositionality of microbiome data and considers the possibility of indirect correlations (ref). Co-occurrence events with correlation of >|0.4| were visualized in Cytoscape v.3.10.0 where nodes represent taxa and edges represent positive and negative co-occurrences according to the SparCC R values. Betweenness and closeness centrality of nodes within the networks were also analyzed in Cytoscape.

### Gene catalog construction and analysis

Protein-coding genes were initially identified using Prodigal v.2.6.3 through the ATLAS workflow (37). Elimination of duplicate genes was achieved through linclust with minid = 0.9 and coverag = 0.9 parameters (38). The quantification of gene abundance per sample was performed using the combine_gene_coverages function within the ATLAS workflow, aligning high-quality filtered reads to the gene catalog via the BBmap suite v.39.01-1 (39). Taxonomic and functional annotations were assigned based on the EggNOG database 5.0, employing eggnog-mapper (v.2.0.1) (40). Subsequently, KEGG annotations were extracted from the output (41-43). Read counts were implemented from the quality control workflow in ATLAS. The resultant outputs from ATLAS were analyzed in RStudio, following the procedures outlined in (https://github.com/metagenome-atlas/Tutorial/blob/master/R/Analyze_genecatalog.Rmd).

To ensure comparability of mapped read fractions for each sample, genes with annotations were acquired and normalized using the median of ratios method through DESeq2 (44). Prefiltering was applied to retain only rows with a count of at least 10 for a minimum number of samples, which was determined based on the number of subjects with high methanogen phenotype. Furthermore, a differential expression analysis was conducted between subjects exhibiting a high methanogen phenotype and other subjects, employing the DESeq2 package in R Studio.

### Gene correlation with taxonomic information

The normalized abundance of genes was then correlated with the CLR transformed abundance of species from the shotgun sequencing data in R. This analysis was performed separately for each age group after the application of ConQuR on the count data. Only the genes of interest coding for enzymes responsible for butyrate and propionate formation were correlated with the taxonomic data using Spearman rank correlations (45). The analysis was plotted in a heatmap using ggplot2.

### Data and software availability

Raw nucleotide data generated and used in the study (Cohorts A and B) can be found in the Sequence Read Archive under the project accession PRJEB72212. All scripts, bracken output, and all the relevant metadata for the cohorts and the concentration of the breath methane (available for cohort A) are provided in (https://github.com/Roxy-mzh/Archaea_And_Aging.

## Results

### Study overview

We studied three distinct cohorts; cohort A, consisting of young adults recruited in Graz, Austria (n=91, ages 19-37 years), cohort B consisting mostly of older adults also recruited in Graz, Austria (n=94, ages 46-86 years) and cohort C with subjects enrolled in the Emilia Romagna (Italy) by Rampelli et al. (19) (n=62, ages 22-109 years) which was geographically very close to the other cohorts and mostly included centenarians (Supplementary Fig. 1).

**Supplementary Fig. 1.**
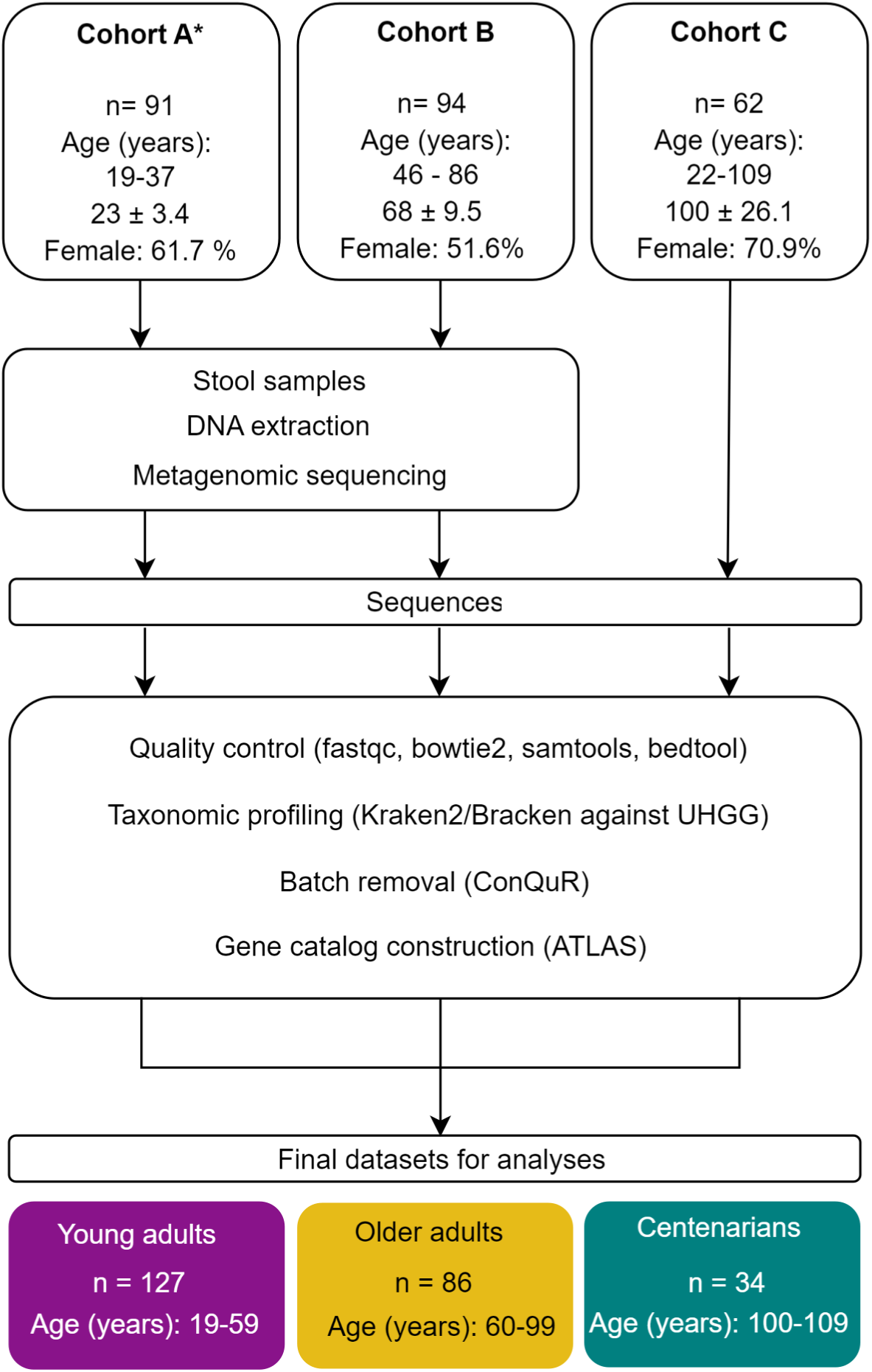
Conceptual outline of study cohorts. Three different study populations were used. For Cohort A, stool samples collected for a study by Kumpitsch et al. (17) were used. Cohorts A and B were collected from the same location but at different time points. Breath methane measurements were recorded for cohort A as indicated by the (*) sign. Samples from Cohorts A and B were processed the same way and a similar method for library preparation was employed. Subjects within Cohort C were enrolled in a study with a close location to cohorts A and B by Rampelli et al. (19), and the deposited sequences were used for further evaluations. In order to mitigate the study effect and remove the bias based on the methods employed in sequencing, ConQuR was used for correcting the read counts.

Overall, the study cohorts included a total of 247 subjects, of three age groups; 127 subjects aged 19–59 y (young adults, “YAs”), 86 subjects aged 60–99 y (older adults, “OAs”), and 34 subjects aged 100-109 (centenarians, “CENT”).

The analysis of age distribution among the three study groups (Cohort A, B, and C) revealed statistically significant differences, as expected. One-way analysis of variance (ANOVA) was conducted to compare the mean ages of participants in each cohort. The results indicated a significant variation in age across the cohorts (F = 391.323, *p* < 0.001), confirming the representation of different age groups by each cohort. On the other hand, a similar distribution of males and females existed within the investigated cohorts as shown by the chi-square test of independence (χ^2^ = 5.871, df = 2, *p* = 0.053).

It is important to note that the three cohorts under study may exhibit variations in numerous potential covariates that have remained unrecorded. Moreover, as differences in sample processing could result in bias in data analysis, we implemented a batch effect correction procedure based on the available metadata (see below).

### Removal of batch effect allows for comparison across datasets

The abundance profiling of the combined datasets based on UHGG showed a total of 4,253 distinct species, which were identified across 247 different samples. In order to remove the batch effects between studies and eradicate the high data variability, we employed the ConQuR. tool.

It was evident that ConQuR significantly diminished the study-related variation observed in the raw count data, as indicated by the Bray-Curtis and Aitchison distance analyses (Supplementary Fig. 2). Although PERMANOVA test showed no difference the Bray-curtis and Aitchison distances before (Bray-curtis *p* = 0.001, Aitchison *p* = 0.001) and after (Bray-curtis *p* = 0.001, Aitchison *p* = 0.001) applying ConQur, the centroids representing the means of the three studies became noticeably closer, and the dispersions and additional characteristics such as the sizes and angles of the ellipses exhibited much greater alignment.

**Supplementary Fig. 2.**
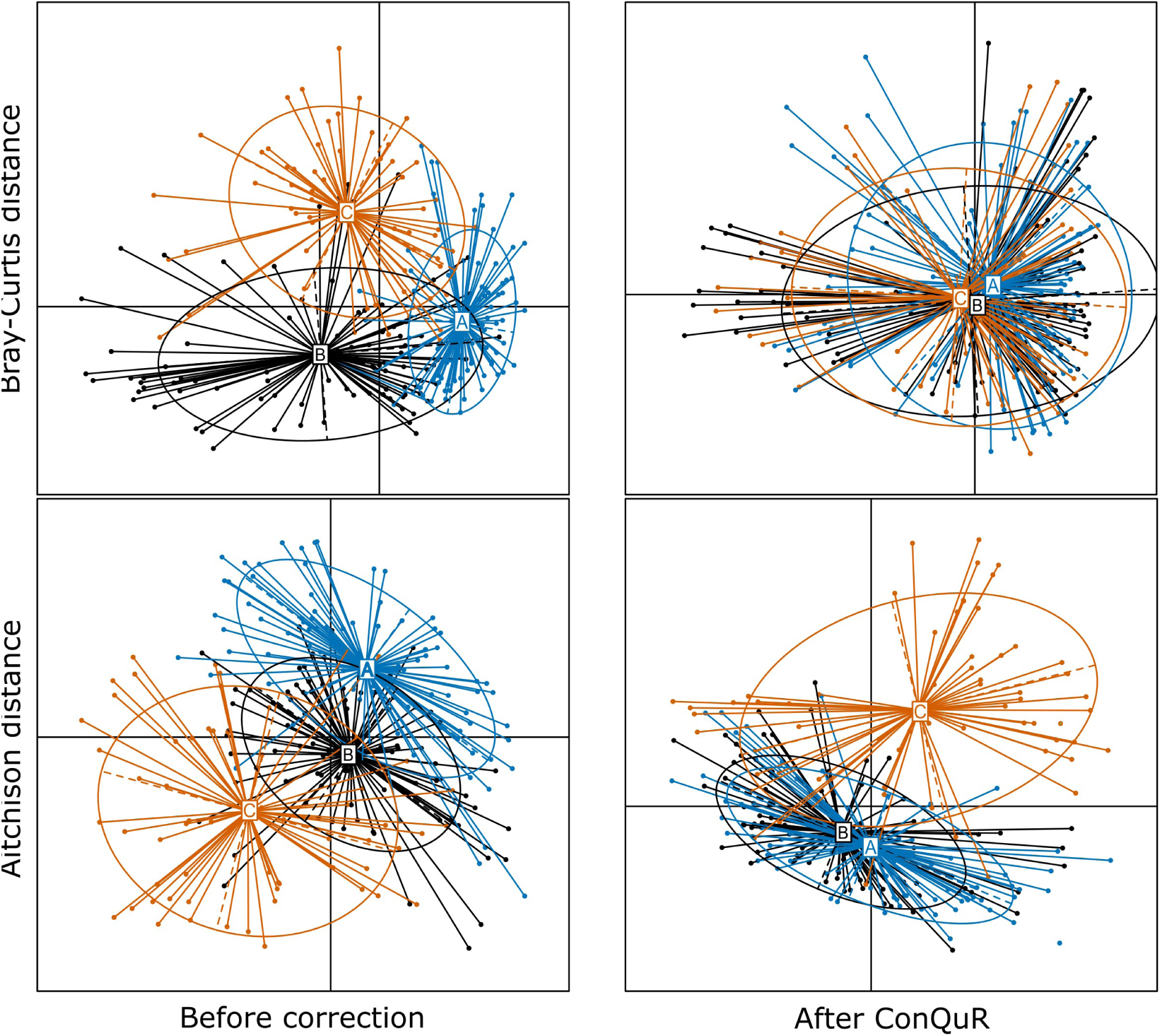
Principal Coordinate Analysis (PCoA) plots were generated to visualize the clustering of study cohorts based on Bray-Curtis and Aitchison dissimilarity computed using raw count data. Each data point on the plot corresponds to a sample, while each ellipse represents a batch (study cohorts A, B, or C), with the centroid denoting the mean. The size of the ellipse reflects the dispersion of data points within each batch, and the angle of the ellipse indicates higher-order characteristics specific to the batch. Furthermore, the ellipse connects the 95th percentile of data points, providing a visual representation of the batch’s overall distribution.

Moreover, based on the PERMANOVA analysis on the count data, the initial R^2^ = 0.10026030 (before applying ConQuR), reduced to R^2^ = 0.01868353 (after applying ConQuR), which was lower than the effect of age (R^2^ = 0.03042544, after ConQuR vs. R^2^ = 0.05358378, before ConQuR) and effect of the presence of high methanogen phenotype (R^2^ = 0.02614169; after ConQuR vs. R^2^ = 0.03092168; before ConQuR).

### Aging is mirrored in the overall microbiome profile

After applying ConQuR for the 247 samples analyzed, the average number of raw reads for metagenomic analyses per subject from each age group ranged from 7,741,637, 6,993,479.8, and 10,103,384 for YAs, OAs, and CENT, respectively, with archaeal reads constituting 0.4972%, 0.4001%, and 0.9476% of all reads corresponding to each age classification in the mentioned order, indicating almost the highest distribution of archaeal read counts in CENT (Mann-Whitney U-test; YAs:OAs *p* = 0.07, YAs:CENT *p* = 0.219, OAs: CENT *p*=0.463) (Supplementary Fig. 3A).

**Supplementary Fig. 3.**
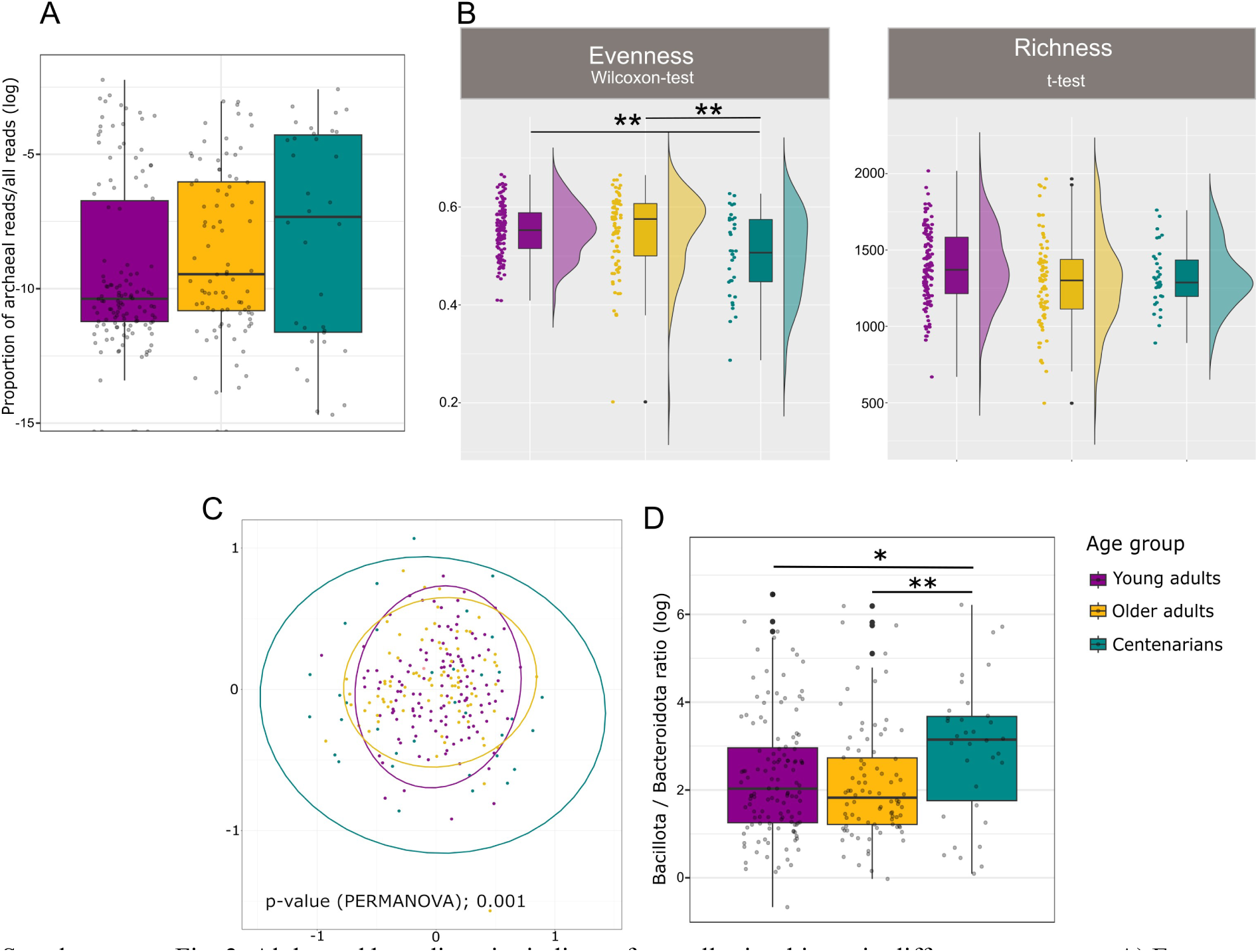
Alpha and beta diversity indices of overall microbiome in different age groups A) Evenness and richness indices tended to decrease with aging. T-test was used for statistical analysis of the richness index due to the normal distribution of the values while evenness values were not normally distributed B) NMDS analysis shows a shift of the clusters based on aging. Statistical significance is indicated by \*\**p* < 0.001\*\**p* < 0.01 and \**p* < 0.05. C). The ratio of Bacillota to Bacteroidota showed an increase with age.

For alpha diversity analysis of the overall microbiome, we calculated the Shannon, richness, and evenness indices. The Shannon index of CENT was significantly lower than that of YAs (Wilcoxon test, *p* = 0.002) and OAs (Wilcoxon test, *p* = 0.005) (Fig. 1A). The same trend was observed for the evenness index as it was significantly lower in CENT compared to YAs (Wilcoxon test, *p* = 0.003) and OAs (Wilcoxon test, *p* = 0.002) (Supplementary Fig. 3B). The richness index decreased with age (Supplementary Fig. 3B), however, this was not significant (Supplementary Fig. 3B). These results suggest a lack of statistical divergence in the alpha diversity of the microbiota community between YAs and OAs; however, alpha diversity of CENT was significantly lower as compared with the other two age groups. This significant drop in the Shannon diversity measure of CENT was in line with observations made in previous studies (12, 46).

**Fig. 1.**
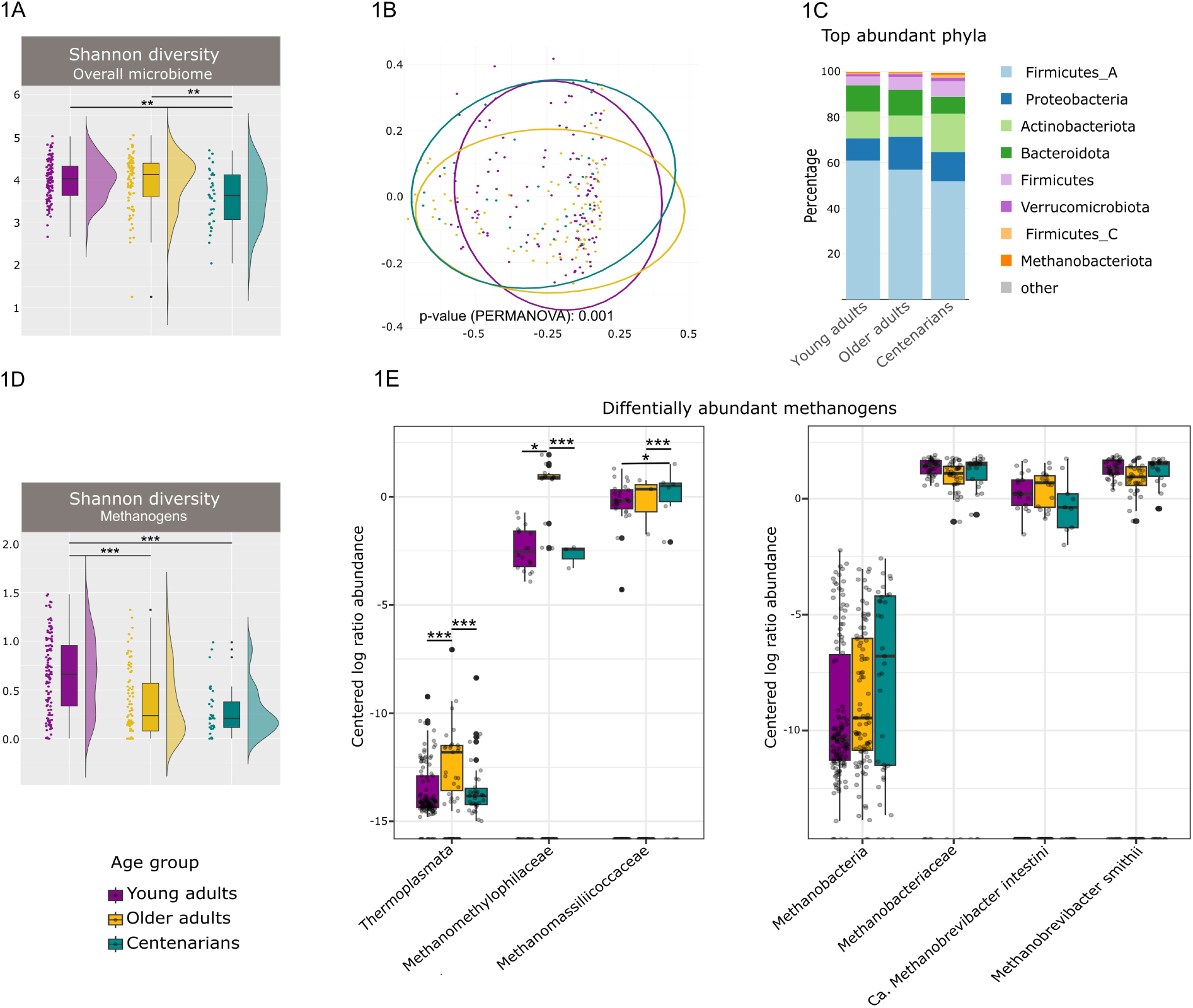
Comparative analysis of fecal microbiome diversity and methanogens in different age groups. (1A) Comparison of Shannon and (1B) beta diversity of fecal microbiomes between YAs, OAs, and CENT. (1C) Stacked bar plot of relative abundances of the top microbial phyla is displayed by age groups (1D) Shannon diversity of methanogens in different age groups. (1E) Box plot of CLR-transformed abundances of the methanogenic archaea in each age group. Line in boxes is a median of index scores, boxes represent interquartile range, whiskers represent lowest and highest values, and dots represent each sample. Statistical significance in 1A and 1D is indicated by \*\**p* < 0.001\*\**p* < 0.01 and \**p* < 0.05. Statistical significance levels of ALDEx2 test in 2E after adjustment for multiple comparison are indicated with \*\*\**q* < 0.001, \*\**q* < 0.01, \**q*< 0.05.

To characterize gut microbial patterns associated with aging, we also performed a β-diversity analysis using principal coordinates analysis (PCoA) and Non-Metric Multidimensional Scaling (NMDS). A major overlap was observed in the PCoA and NMDS plots, however, PERMANOVA under 999 permutations showed a significant difference (*p* < 0.001; stress (NMDS): 0.2577) between the beta-diversity of the three age groups (Fig. 1B, Supplementary Fig. 3C), as observed previously (6).

With respect to the bacteriome, we could observe that at phylum level, Firmicutes, Proteobacteria, Actinobacteriota, and Bacteroidota were the dominant bacterial taxa in each age group, which was in accordance with previous reports with different cohorts (Fig. 1C) (47). However, both Firmicute_A and Bacteroidota exhibited significantly lower levels in CENT when contrasted with YAs and OAs (Firmicute_A *p* < 0.01, *q* < 0.01; Bacteroidota *p* < 0.001, *q* < 0.001; Wilcoxon test). On the other hand, the ratio of whole Bacillota (former: Firmicute) phylum (including Firmicutes, Firmicute_A, Firmicute_B, and Firmicute_C) to Bacteroidota, which is commonly used as an index for the composition of the gut microbiome showed significant elevation in CENT (ratio ≈ 56.83065) compared to YAs (ratio ≈ 31.76721) and OAs (ratio ≈ 28.26067) (Mann-Whitney U-test, YAs:CENT *p* = 0.02, OAs: CENT *p* = 0.006, YAs:OAs *p* = 0.367) (Supplementary Fig. 3D) as described previously (48).

### The composition of methanogens in centenarians is similar to that of young adults

The potential impact of archaea, as the understudied members of the human microbiome, on aging was investigated in more detail. Aging affected the alpha diversity of these microbial members as exhibited by the statistically significant (*p* < 0.001) decrease in the Shannon index of methanogenic species across the various age groups under investigation (Fig. 1D), which was in contrast with previous reports based on 16S rRNA gene amplicon sequencing (15).

Age-associated changes in the archaeome profile was not manifested on phylum level, but were observed on class, order, family, and species levels. The class Methanobacteria exhibited an observable elevation in abundance in relation to the aging process, however, this increase was not significant (YAs: OAs *p* = 0.025, *q* = 0.162, YAs: CENT *p* = 0.367, *q* = 0.489; ALDEx). On the other hand, the class Thermoplasmata showed a statistically significant increase of relative abundance in OAs in comparison to both YAs (*p* < 0.001, *q* < 0.001) and CENT (*p* < 0.001, *q* = 0.00102) (Fig. 1E, Supplementary Table).

Further exploration at the family taxonomic level revealed that this significant increase of Thermoplasmata in OAs is mostly driven by the Methanomethylophylaceae family (OAs:YAs *p* < 0.001, *q* < 0.001, OAs:CENT *p* < 0.001, *q* < 0.001; Fig. 1E). Similar elevation trends were also observed for Methanomassiliicocaceae as another member of the Thermoplasmata class. However, this elevation was observed not only in OAs, but also in CENT (OAs:YAs *p* > 0.5, *q* > 0.5, CENT:YAs *p* = 0.005, *q* = 0.019, CENT:OAs *p* < 0.001, *q* = 0.0013; Fig. 1E, (Supplementary Table). This observed increased abundance of Methanomassiliicocaceae is consistent with previous reports (16, 49). Interestingly, Methanobacteriaceae showed decreased abundance levels in OAs compared to the other age groups, however, this decrease was not statistically significant (OAs: YAs *p* = 0.0102, *q* = 0.062; OAs:CENT *p* > 0.5, *q* > 0.5; Fig. 1E, Supplementary Table)

An interesting dynamic was observed for *Methanobrevibacter* species, indicating an important transition of the archaeome in OAs. The predominant gut archaeal species, *Methanobrevibacter smithii* exhibited a decreased abundance in OAs relative to both YAs and CENT (*p* > 0.5, *q* > 0.5). Conversely, *Ca.* M. intestini, the newly separated clade within the *Methanobrevibacter* genus (18), displayed an increase in abundance among OAs compared to YAs (*p* = 0.064, *q* = 0.263), while its abundance dropped in CENT in comparison (CENT: YAs *p* = 0.024, *q* = 0.0905, CENT: OAs *p* > 0, *q* > 0.5) (Fig. 1E, Supplementary Table)

In short, our observations show that while there is a decrease in the variety of methanogens in both OAs and CENT, the composition of methanogenic archaea found in CENT is more similar to those in YAs than in OAs. This similarity is especially noticeable at archaeal family and species levels, where we see similar patterns in the abundance of Methanomethylophylaceae and Methanobacteriaceae (at family level) as well as *M. smithii* (at species level) in YAs and CENT. On the other hand, OAs have a higher abundance of *Ca.* M. intestini, which appears with a diminished abundance in CENT. This suggests that *Ca*. M. intestini may play a substantial role during transition to longevity.

### The two predominant *Methanobrevibacter* species co-exist and demonstrate co-occurrence with health-associated bacterial species

Microbial networks, which are constructed based on correlations in species abundances, offer insights into co-occurrences among microbes within a community. In order to gain a better understanding of the microbes with potential co-occurrence with *M. smithii* and *Ca.* M. intestini in the gut across different age groups, we utilized abundance data of taxa from each age group to generate three distinct networks (Fig. 2A). Of note, co-occurrence in this correlation-based network does not indicate physical or biochemical interactions among microbes, but rather is indicative of niche sharing (50).

**Fig. 2.**
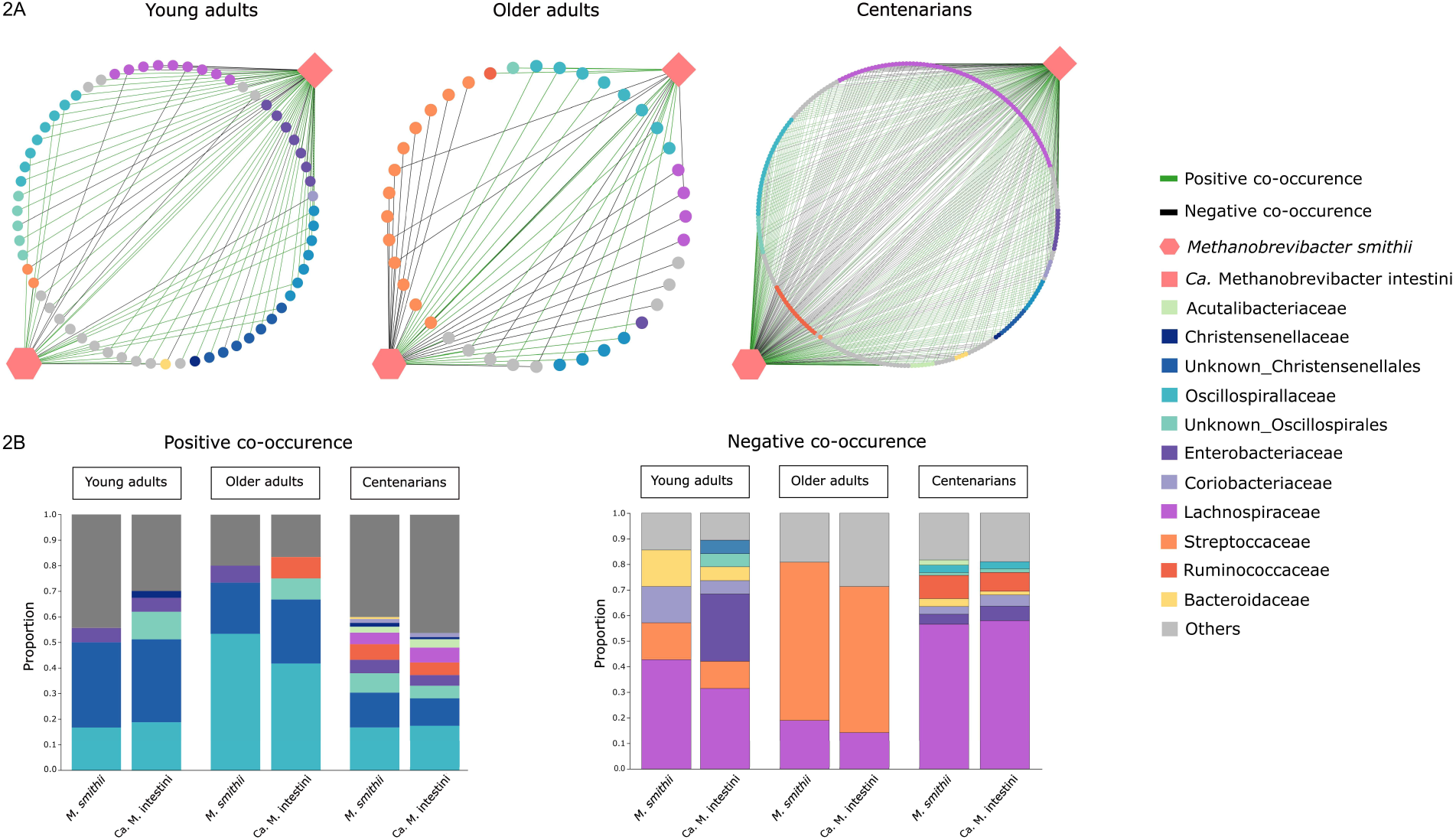
SparCC co-occurrence networks of *M. smithii* and *Ca.* M. intestini in all samples irrespective of the presence of high methanogen phenotype (2A). Each node represents a single microbial species, and each edge a single association between a pair of microbial species. Positive and negative SparCC co-occurrences are indicated in green and black, respectively. These co-occurrences are elaborated in (2B).

Looking closer at the co-occurrence networks, CENT demonstrated the highest number of node degrees for both *M. smithii* and *Ca.* M. intestini (Table 1), indicating a high interconnectivity for both *Methanobrevibacter* species in particular in CENT microbiomes, while in OAs this interconnectivity was the lowest. These findings confirmed that microbial networks of *Ca.* M. intestini and *M. smithii* indeed undergo changes throughout human aging.

**Table 1.** Parameters of *M. smithii* and *Ca.* M. intestini networks with the gut microbial community of the studied subjects.

| Archaeal species | Network parameter | Network inference for all studied subjects |  |  | Network inference for subjects with high methanogen phenotype |  |  |
| --- | --- | --- | --- | --- | --- | --- | --- |
|  |  | YAs | OAs | CENT | YAs | OAs | CENT |
| *M. smithii* | Node degree (+/-)* | 25 (18+, 7-) | 37 (15+, 21-) | 231 (132+, 98-) | 82 (46 - /36 +) | 117 (62+/55-) | 169 (47-, 122+) |
|  | Betweenness centrality | 0.246 | 0.832 | 0.659 | 0.507 | 0.902 | 0.790 |
|  | Closeness centrality | 0.627 | 0.911 | 0.871 | 0.442 | 0.678 | 0.734 |
| *Ca. M. intestini* | Node degree (+/-)* | 56 (37+, 19-) | 19 (12+, 7-) | 190 (121+, 69-) | 156 (90 -/66+) | 48 (27+/21-) | 120 (60 +/60 -) |
|  | Betweenness centrality | 0.876 | 0.251 | 0.424 | 0.864 | 0.416 | 0.593 |
|  | Closeness centrality | 0.901 | 0.661 | 0.770 | 0.626 | 0.421 | 0.649 |

\* Positive (co-presence) /negative (mutually excluded) association

Based on betweenness and closeness centrality measures as well as node degree values (Table 1), *Ca.* M. intestini was the keystone species in driving the network with other microbes in YAs, while in OAs and CENT, *M. smithii* was the major driver of the network (Table 1). These results suggest that *Ca.* M. intestini might pave the way for the prominence of *M. smithii* in the overal function and stability of the microbial network later on with age.

By taking a closer look at the various microbial species within the co-occurrence network of these two predominant methanogens we delved into their role in shaping the assembly of microbial communities with *M. smithii* and *Ca.* M. intestini. Specifically, our observations indicate a consistent co-occurrence of *M. smithii* and *Ca.* M. intestini across all three age groups of YAs, OAs, and CENT (Fig. 2A). This finding is somewhat surprising, as both archaea require similar ressources for their growth and competition-based suppression of one species would appear logical.

Furthermore, our results highlight the prevalence of certain taxa within these association networks of *M. smithii* and *Ca.* M. intestini across all age groups. Notably, Oscillospirales/Oscillospiraceae as well as Christensenellales/Christensenellaceae predominantly signify co-presence and niche-sharing with these two archaeal species in all three age groups (Fig. 2A and Fig. 2B). The positive co-occurrence of *Oscillospira spp.* and members of Christensenellaceae with *M. smithii* has been also documented before (17, 51).

In contrast, although some members of Lachnospiraceae, such as *Roseburia hominis*, *Blautia hansenii*, *Blautia massiliensis* showed positive co-occurrences with these two archaeal species in CENT, a high proportion of this bacterial family was predominantly associated with negative co-occurrences (co-absence or mutual exclusion) in all three age groups (Fig. 2C).

It is worth noting that certain abundant co-occurring bacterial taxa with *M. smithii* and *Ca.* M. intestini exhibited an age-specific pattern. For instance, *Streptococcus* emerged as the dominant taxon showing mutual exclusion association with *M. smithii* and *Ca.* M. intestini mostly in OAs (Fig. 2C). It shall be noted that an increase of *Streptococcus* species could indicate an oralisation of the gastrointestinal microbiome e.g. by the use of proton pump inhibitors (52).

Moreover, although some positive co-occurrences of *M. smithii* and *Ca.* M. intestini were observed with opportunistic pathogens within the family Enterobacteriaceae, mutual exclusion with pathogens (*Klebsiella*, *Salmonella* and *Proteus*) were mostly negative, in YAs and CENT.

### High methanogen phenotypes are twice as common in centenarians

Since the focus of our study was on the human archaeome and its impact on healthy aging, a phenotypic grouping of individuals according to their archaeal profile was necessary. The most easily measurable physiological effect of high intestinal archaeal load is the exhalation of substantial amounts of methane (17).

For cohort A, methane breath content information was available and we confirmed a statistically significant association between the manifestation of positive breath methane production (≥5 ppm) and the abundance of archaea (Mann-Whitney U, *p* < 0.001) and, in particular, *Methanobrevibacter* signatures (Mann-Whitney U, *p* < 0.001). Interestingly, our results indicated that both *M. smithii* and *Ca.* M. intestini contributed to methane production (Supplementary Fig. 4A).

**Supplementary Fig. 4.**
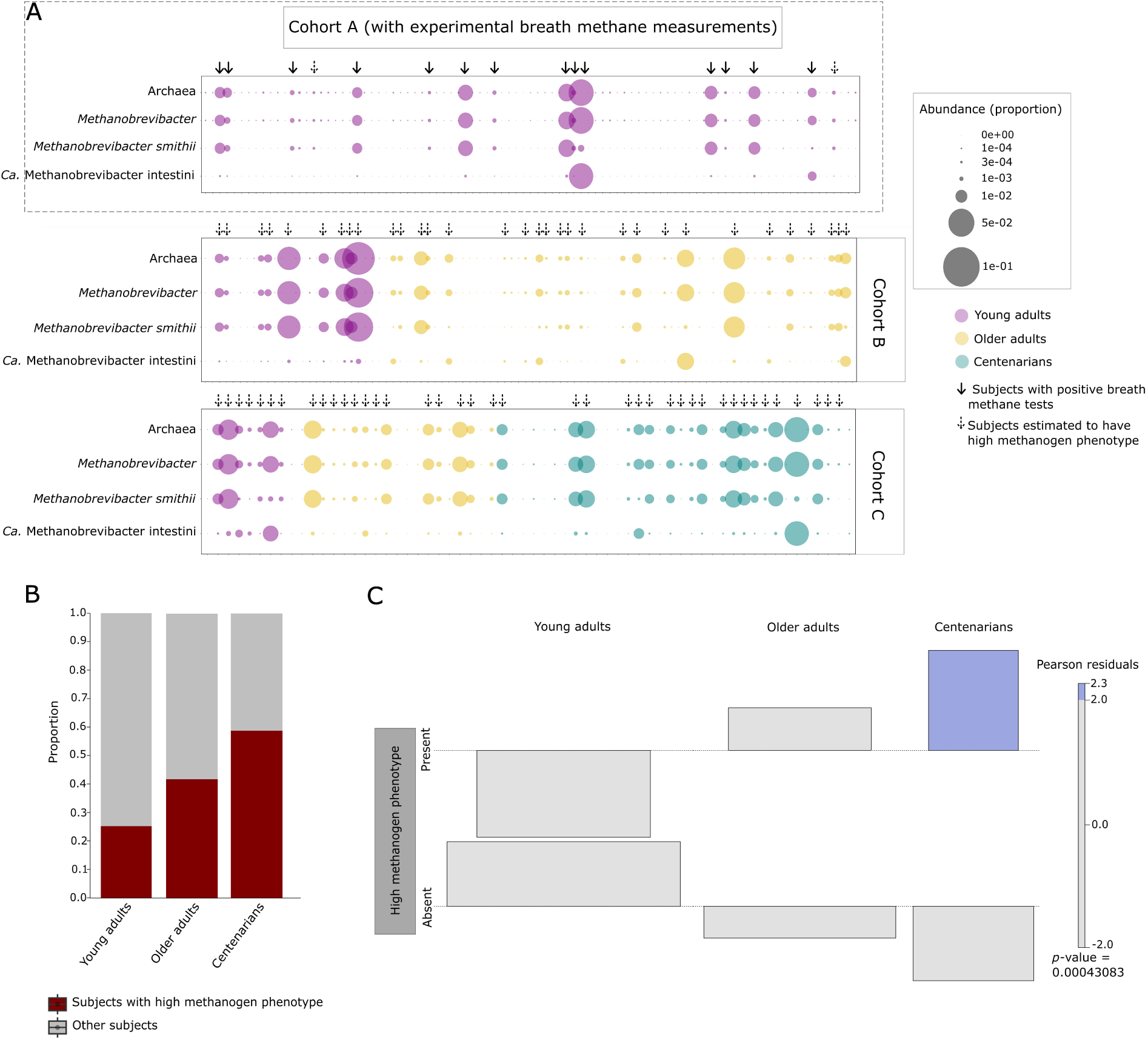
Age-dependent prevalence of high methanogen phenotype. A) Methane production in cohorts, with full arrows indicating samples showing >5 ppm breath methane. Some samples had high *Methanobrevibacter* abundance despite negative breath test results. Samples estimated to have high methanogen phenotype are shown with dashed arrows. B) The prevalence of high methanogen phenotype increases with age. C) Association plot visualizing that high frequency of high methanogen phenotype is associated with the CENT age group rather than other age groups. Area of the box is proportional to the difference in observed and expected frequencies of the presence of high methanogen phenotype. The baseline (dotted line) indicates independence of high methanogen phenotype to aging. The boxes rising above the baseline indicate that the observed frequency of a cell is greater than the expected one (if the data were random), and *vice versa*. Cells representing negative residuals are drawn below the baseline and vice versa. The width of each of the bar elements in the mosaic reflects the relative magnitude of its value.

A bimodal pattern has been demonstrated for the prevalence of *Methanobrevibacter* in the human gut, indicating that it is either highly prevalent or almost absent (53). However, some subjects might still show a certain abundance of *Methanobrevibacter,* while not being categorized as methane emitters. According to our observations *Methanobrevibacter* abundance of ≥ 0.03% could be considered as a threshold for subjects with positive (> 5 ppm) breath methane. Despite yielding negative results in breath test measurements within cohort A, two samples (Sample IDs: ORS_116 and ORS_94) exhibited a significant abundance of *Methanobrevibacter* at levels akin to those observed in individuals displaying elevated breath methane levels (indicated by dashed arrows in Supplementary Fig. 4A). This could potentially be attributed to technical inaccuracies in sample processing and reads-outs or physiological peculiarities at this time point. Utilizing this established threshold, we then conducted an evaluation to define the subjects with high methanogen phenotype within cohorts B and C (Supplementary Fig. 4A).

The prevalence of the subjects with high methanogen phenotype within distinct age groups was as follows: 25.2% (32/127; 95% CI: 19.6-31.6%) among YAs, 41.86% (36/86; 95% CI: 32.6-51.1%) among OAs, and 58.82% (20/34; 95% CI: 41.3-73.1%) among CENT (Supplementary Fig. 4B). This observation suggests an elevated prevalence of subjects with high methanogen phenotype with respect to aging. According to the results of Pearson’s chi-square test the prevalence percentages of subjects with high methanogen phenotype appear to be significantly different among different age groups (chi-square = 13.762, df = 2, *p* = 0.001027), and the presence of high methanogen phenotype and age group variables are statistically significantly associated (chi-squared = 15.458, df = 2, *p* < 0.001) (Supplementary Fig. 4C). In fact, based on these results, it was evident that the highest association between the presence or the frequency of high methanogen phenotype was in centenarians, while the reduced frequency of high methanogen phenotype was positively associated with YAs.

### The presence of high methanogen phenotype affects microbiome characteristics across age groups

A significantly higher Shannon diversity was observed in subjects with high methanogen phenotype compared to other subjects in YAs and OAs (Fig. 3A) (Shannon index = 4.247 ± 0.353, *p* < 0.001 for YAs; Shannon index = 4.171 ± 0.265, *p* = 0.025 for OAs) (consistent with previous results (17)). There was also a consistent and statistically significant elevation in microbial richness attributed to the high methanogen phenotype, across all three age groups (*p* < 0.001; t-test), indicating a higher diversity of microbial signatures in the presence of high methanogen phenotype. The evenness measure, however, exhibited significant elevation with respect to the high methanogen phenotype only among YAs (*p* = 0.002; t-test) (Supplementary Fig. 5A).

**Fig. 3.**
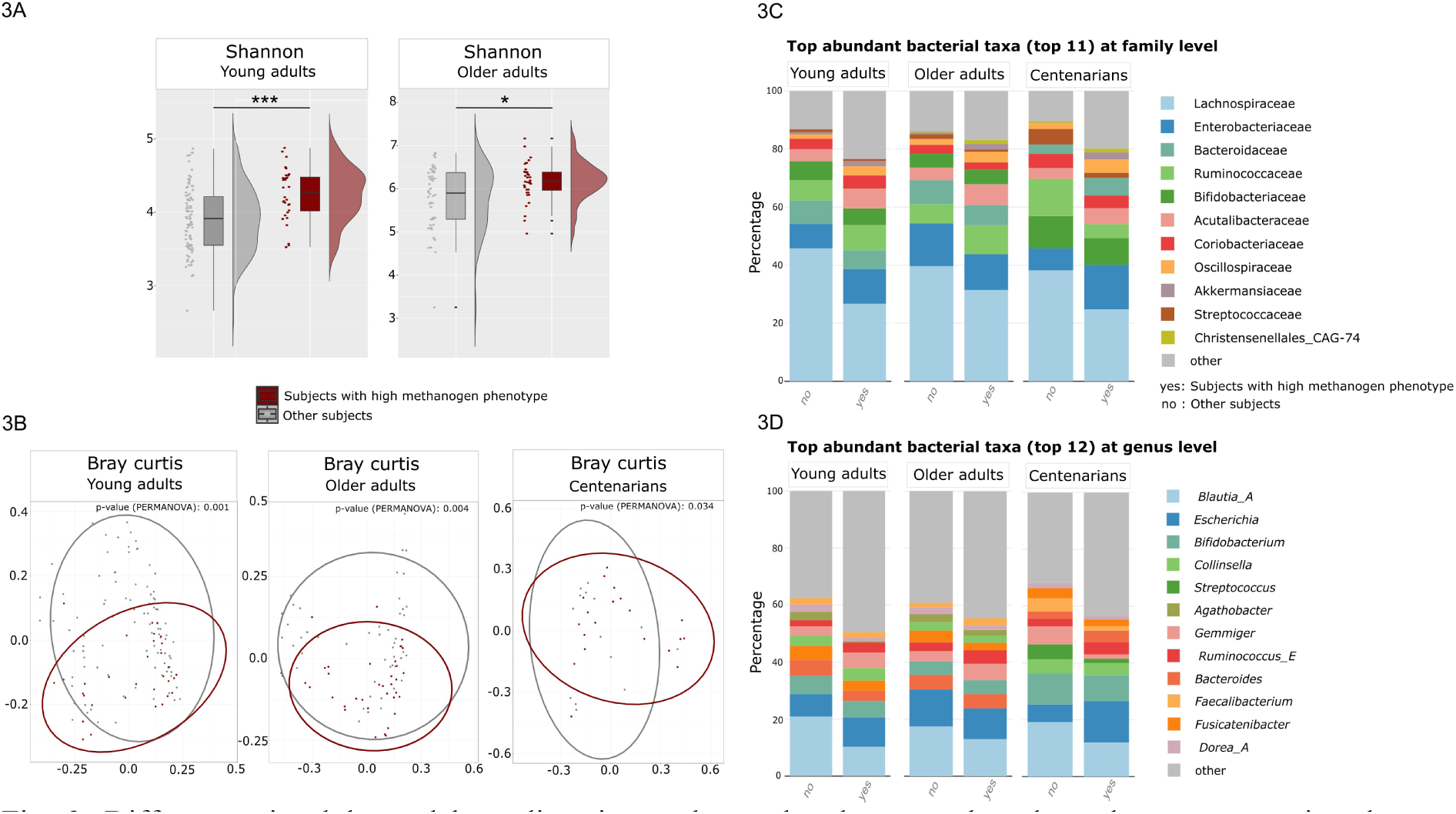
Differences in alpha and beta diversity, and top abundant taxa based on the metagenomics shotgun sequencing between subjects with high methanogen phenotype and other subjects in different age groups. A) An examination of Shannon diversity index revealed significant differences in alpha diversity based on high methanogen phenotype in both YAs and OAs (***p< 0.001, *p< 0.05). However, all subjects either with or without high methanogen phenotype within CENT showed similar trends regarding Shannon index. B) The microbiome of subjects with the high methanogen phenotype clustered significantly differently in the PCoA plots, regardless of age classification. However, this significance was notably higher only in YAs and OAs. C) Bar chart of the most abundant bacterial taxa at family level and D) genus level compared regarding the presence of high methanogen phenotype with respect to each age classification of YAs and OAs, as well as CENT.

**Supplementary Fig. 5.**
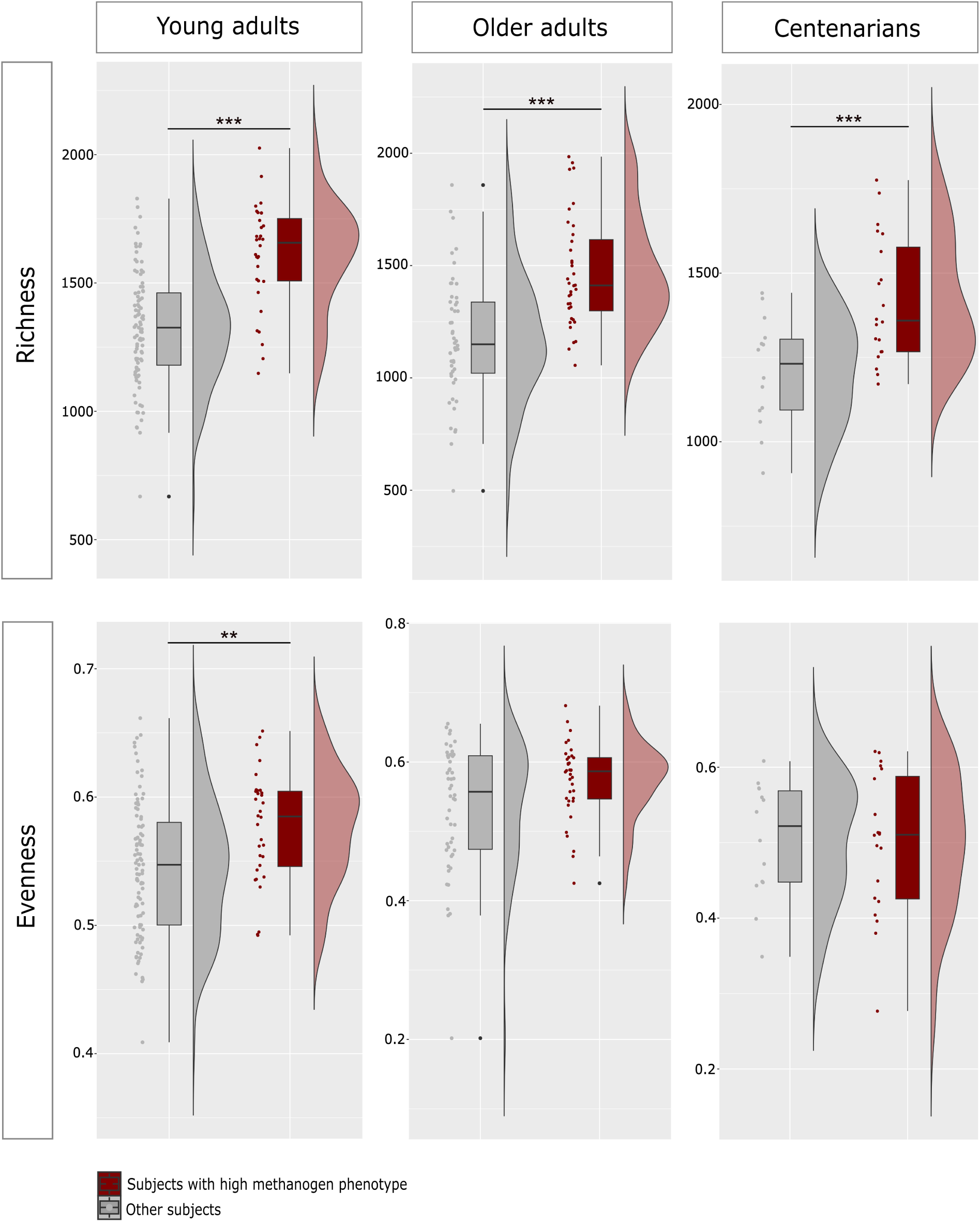
An examination of the richness index revealed significant differences based on the presence of high methanogen phenotype irrespective of the age classification, with those with high methanogen phenotype showing significantly higher richness. However, the evenness index was only significantly higher in the presence of high methanogen phenotype in YAs. ***p< 0.001, **p< 0.01.

The microbial community composition revealed statistically significant differentiation between clusters of subjects characterized by the presence of the high methanogen phenotype and those lacking it (within the age groups of YAs *p* = 0.001, OAs *p* = 0.004, and CENT *p* = 0.034) (Fig. 3B).

It is important to note that the variability within the microbial communities of subjects with high methanogen phenotype was notably higher in both YAs and OAs compared to CENT (YAs *p* < 0.001, OAs *p* = 0.005, CENT *p* = 0.028; PERMANOVA). The finding that subjects with high methanogen phenotype in CENT do not exhibit a strongly significant difference in their microbiome compared to those subjects of YAs and OAs might suggest that the gut microbiota of subjects with high methanogen phenotype within the CENT might possess less unique microbiota as compared with other subjects in this age group.

The analysis results centered around the presence of the high methanogen phenotype revealed consistent variations in the abundance of some specific taxa, regardless of their categorization within specific age groups. Specifically, the family Lachnospiraceae and at genus level *Agathobacter, Blautia, Dorea* (known butyrate-producing taxa within the family Lachnospiraceae (54-56), as well as *Bacteroides* were reduced in relative abundance in subjects with high methanogen phenotype across all age groups (Fig. 3B, Fig. 3C, Supplementary Fig. 5A, Supplementary Fig. 5B, Supplementary Table). Additionally, diminished abundance of the family Streptococcaceae and the genus *Streptococcus* was documented in these subjects across all age groups (Fig. 3C, Supplementary Fig. 6A, Supplementary Fig. 6B), while Acutalibacteriaceae, Christensenellales_CAG-74, and Oscillospiraceae showed high abundances within the high methanogen phenotype subjects regardless of their age (Fig. 3C, Supplementary Fig. 6A, Supplementary Fig. 6B, Supplementary Table).

**Supplementary Fig. 6.**
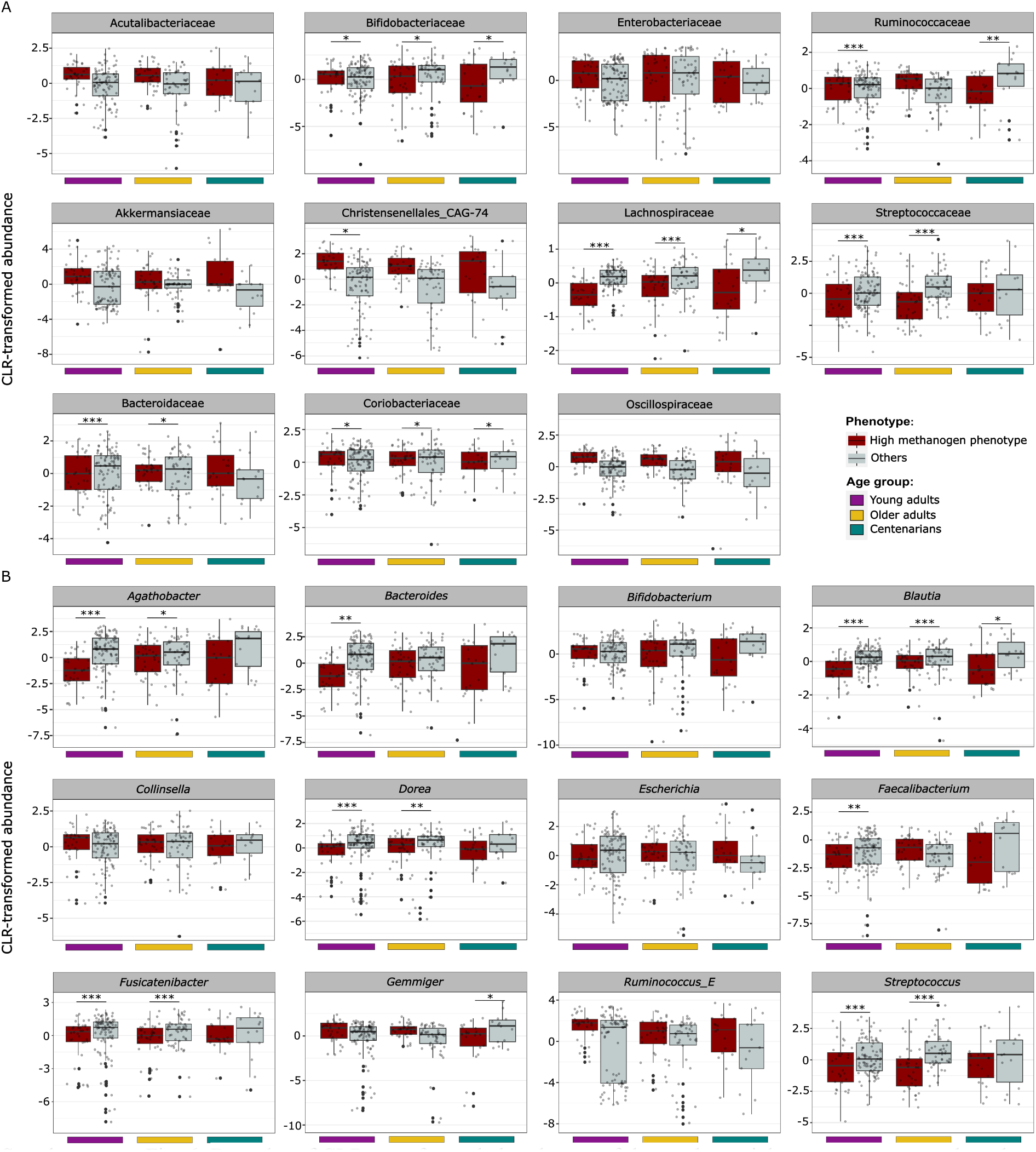
Box plot of CLR-transformed abundances of the top bacterial taxa per age group based on the presence of high methanogen phenotype. A) Top bacterial taxa at family level per age group. B) Top bacterial taxa at genus level. Significance levels are indicated as ***q < 0.001, **q < 0.01, *q< 0.05, for differentially abundance testing by ALDEx.

The microbial composition of those with a high methanogen phenotype was particularly similar in YAs and OAs. In contrast, CENT exhibited slightly varied microbial composition for the high methanogen phenotype, highlighting a distinct microbial ecosystem within this age group. Particularly, a noteworthy observation was made within CENT genera *Gemmiger* and *Faecalibacterium* as well as Ruminococcaceae demonstrated a marked reduction in subjects with the high methanogen phenotype (Fig. 3C, Fig. 3D, Supplementary Fig. 6A, Supplementary Fig. 6B, Supplementary Table). On the other hand, Ruminococcaceae showed increased abundance in subjects with high methanogen phenotype in YAs and OAs. Interestingly, although not statistically significant, *Ruminococcus_E* was highly abundant in subjects with high methanogen phenotype within all three age groups (Fig. 3D, Supplementary Fig. 6B, Supplementary Table), which was consistent with previous reports (17). *Ruminococcus* demonstrates a significant correlation with dietary fibers, owing to its efficient breakdown of microcrystalline cellulose. Previous studies conducted by researchers have indeed elucidated a link between cellulose degradation and the subsequent emission of methane (57).

### Co-occurring bacterial consortium of predominant *Methanobrevibacter* spp. is complex and dynamic in subjects with high methanogen phenotype across all age groups

When examining the microbial networks of *M. smithii* and *Ca.* M. intestini in individuals exhibiting a high methanogen phenotype, it became apparent that the stability of the microbial networks for these two archaeal species remained consistent in these subjects, irrespective of age categorization. Each network displayed comparable complexity. Interestingly, the network complexity and co-occurring microbes were mostly similar in subjects with high methanogen phenotype within YAs and CENT. The microbiome composition and co-occurring taxa are often linked to health status. In aging, OAs commonly experience inflammaging, a chronic low-grade inflammation where the microbiome plays a pivotal role (58). Centenarians may exhibit a unique microbial profile associated with superior health and longevity, reflecting a more similar composition of co-occurring taxa.

The betweenness and closeness centrality metrics for *M. smithii* and *Ca.* M. intestini exhibited comparable patterns to those seen in all subjects, irrespective of the presence of high methanogen phenotype (Table 1). Among YAs with high methanogen phenotype, *Ca.* M. intestini played a central role in the microbial network, whereas in OAs and CENT with high methanogen phenotype, *M. smithii* emerged as the primary driver of the microbial network (Table 1).

Taking a closer look at *M. smithii* and *Ca.* M. intestini edges within networks, mutual exclusion of these archaeal species with members of Ruminococcaceae, Bacteroidaceae, and Streptococcaceae/Streptococcus, and co-presence with Oscillospirales/Oscillospiraceae as well as Christensenellales were observed (Supplementary Fig. 7), which was similar to the trends observed before irrespective of the presence or absence of high methanogen phenotype.

**Supplementary Fig. 7.**
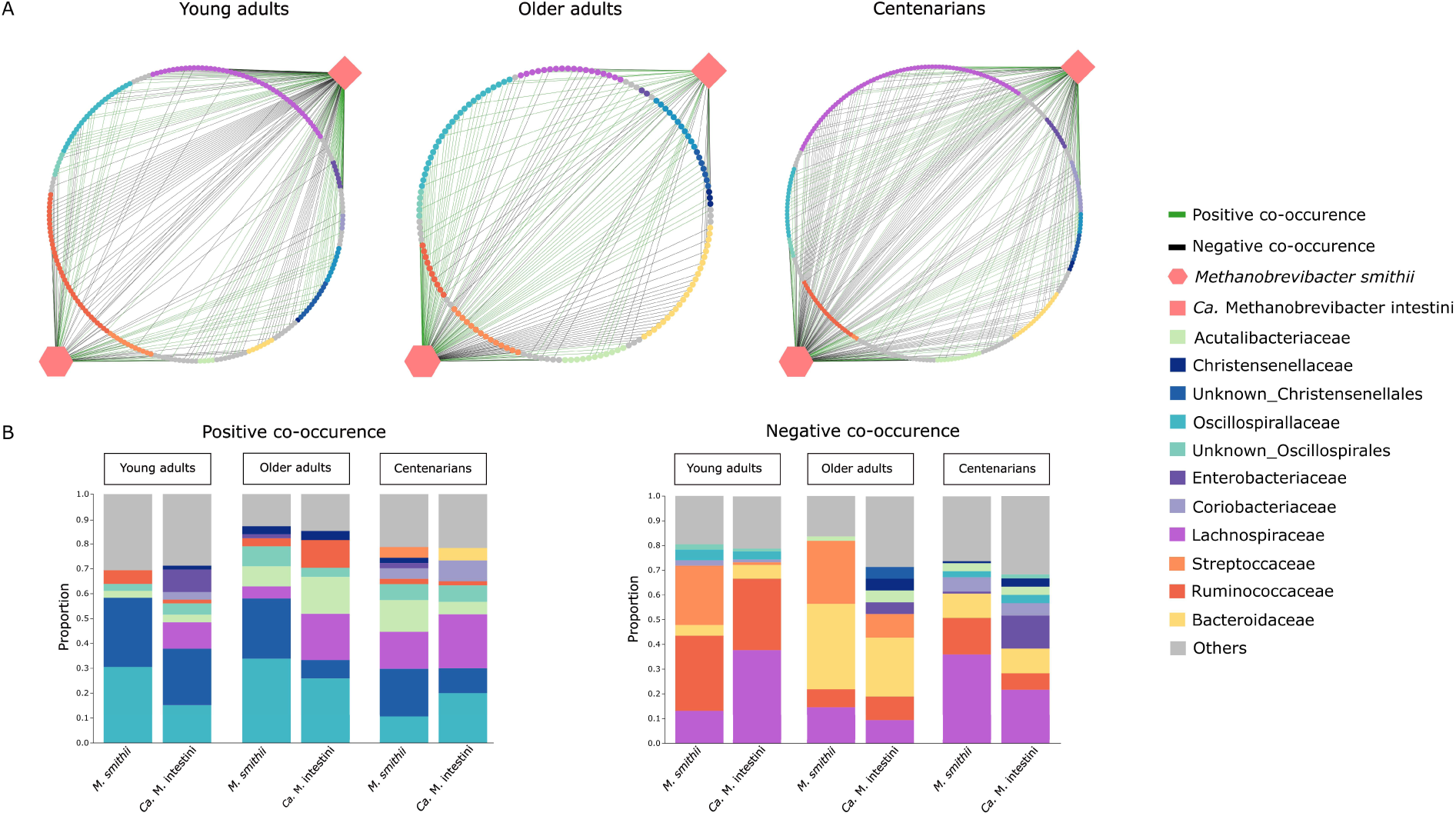
SparCC co-occurrence networks of *M. smithii* and *Ca.* M. intestini in samples with high methanogen phenotype in different age groups of YAs, OAs, and CENT (2A). Positive and negative SparCC co-occurrences are indicated in green and black, respectively. The details of these co-occurrences are shown in more detail in (2B).

Of note, while most members of Lachnospiraceae, including *Acetatifactor* (YAs, CENT), *Agathobacter* (all age groups), *Blautia* (all age groups), *Dorea* (all age groups), *Eubacterium* (YAs, CENT), *Fusicatenibacter* (all age groups), *Lachnospira* (YAs, CENT), and *Roseburia* (YAs, CENT), showed co-presence with *M. smithii* and *Ca.* M. intestini, some of the genera within this family including *Eisenbergiella* (all age groups) and *Mediterranibacter* (all age groups), showed mutual exclusion with these archaeal species. This could be attributed to potential different ecological roles or metabolic functions of these bacteria leading to competition or niche differentiation as well as difference in their adaptation to specific environmental conditions.

### High methanogen phenotype is associated with the upregulation of genes involved in butyrate and propionate production

The decrease in Lachnospiraceae, recognized as butyrate-producing bacteria, has been documented in CENT in several studies, irrespective of the geographical region (13, 59, 60).

This prompts the hypothesis that individuals having detectable *M. smithii* and *Ca.* M. intestini in their gut microbiome may better cope with the decline of these bacteria during aging. This is suggested by the consistent co-occurrence of these archaeal species with Oscillospiraceae, another known butyrate-producing component of the gut microbiota. Moreover, an increased abundance of Oscillospiraceae was observed in subjects exhibiting a high methanogen phenotype (Fig. 2, Supplementary Fig. 6, Supplementary Fig. 7).

Diverse and abundant genes related to butyrate metabolism are present in the metagenomic dataset. Specifically, pathways such as the butyryl-CoA:acetate-CoA pathway and the butyrate kinase pathway contribute to the formation of butyrate. The abundance of genes encoding enzymes directly involved in butyrate production was analyzed. Interestingly, numerous genes related to butyrate metabolism exhibited varying levels of abundance in the metagenome datasets between individuals with high methanogen phenotype and others (Fig. 4A).

**Fig. 4.**
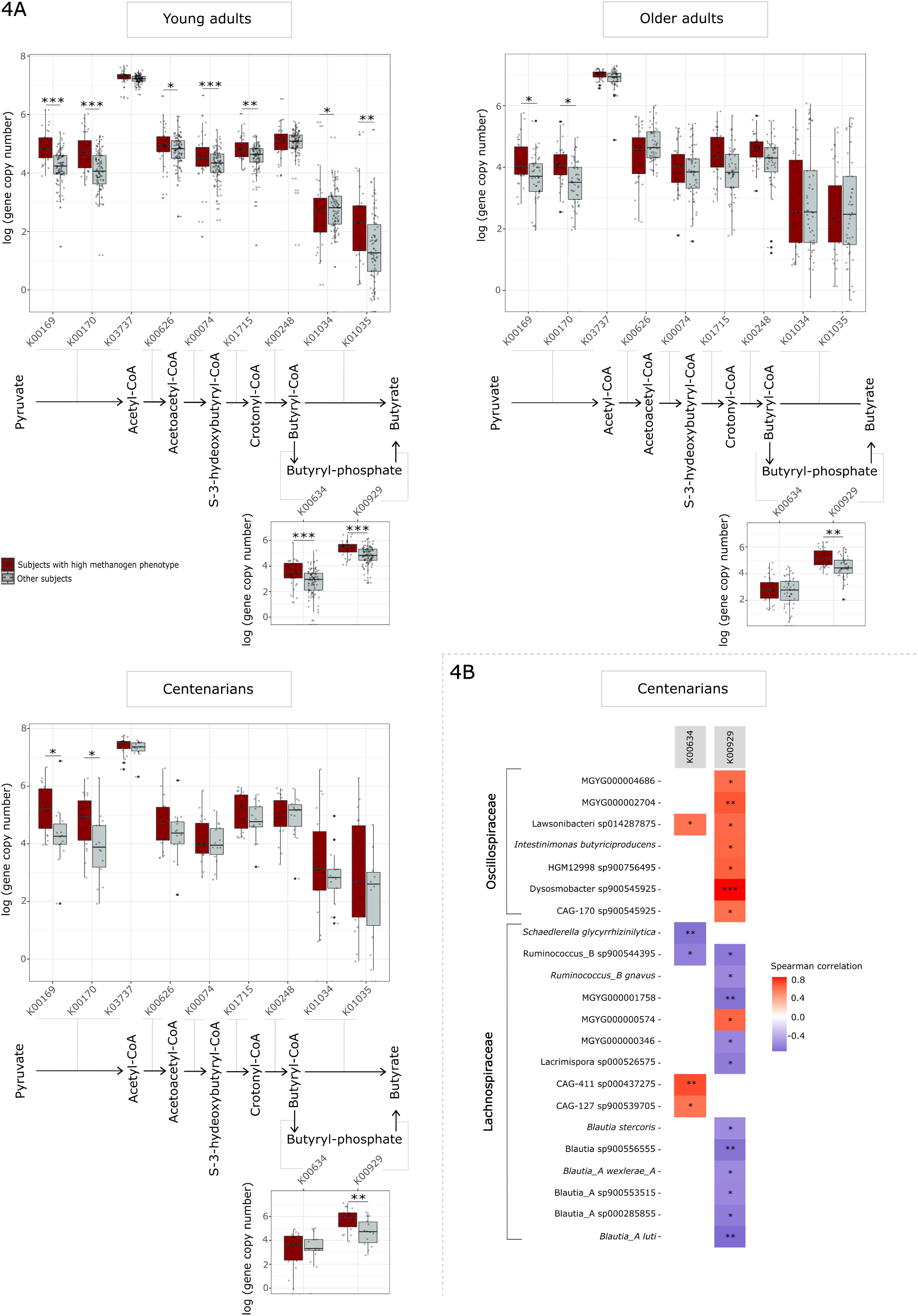
A) Boxplot of gene copy numbers (metagenomic abundance) involved in butyrate production. K00169, K0017, K03737: Pyruvate ferredoxin oxidoreductase [EC: 1.2.7.1]; K00626: Acetyl-CoA acetyltransferase [EC: 2.3.1.9]; K00074: 3-hydroxylbutyryl-CoA dehydrogenase [EC: 1.1.1.157]; K01715: Enoyl-CoA hydratase [EC: 4.2.1.17]; K00248: butyryl-CoA dehydrogenase [EC: 1.3.8.1]; K01034, K01035: Acetate-CoA transferase [EC: 2.8.3.8]; K00634: phosphate butyryltransferase [EC: 2.3.1.19]; K00929: butyrate kinase [EC: 2.7.2.7]. Significance levels are indicated as ***q<0.001, **q<0.01, *q<0.05, for differentially abundance testing by DESeq2. B) Correlation of butyrate kinase pathway genes in centenarians with bacterial taxa. ***q<0.001, **q<0.01, *q<0.05.

In YAs, the majority of genes that play a role in butyrate production, involving both the butyryl-CoA:acetate-CoA and butyrate kinase pathways, exhibited notably higher levels in individuals exhibiting a high methanogen phenotype compared to other subjects (Fig. 4A). One exception was K03737 that is responsible for encoding pyruvate ferredoxin oxidoreductase which, while displaying a tendency toward increased expression, almost reached statistical significance (p = 0.025, q = 0.054).

In OAs and CENT, there was a significant elevated level of genes associated with pyruvate ferredoxin oxidoreductase (K00169, K00170) within the butyryl-CoA:acetate-CoA pathway (q < 0.05) (Supplementary Table). Although not achieving statistical significance, other genes in this pathway displayed a trend of increased expression in these subjects. Intriguingly, within these age groups, butyrate kinase (K00929), the terminal enzyme in the butyrate kinase pathway responsible for synthesizing butyrate, exhibited statistically elevated levels (q < 0.01). Additionally, phosphate butyryltransferase (K00634) displayed an increasing trend in subjects with a high methanogen phenotype, indicating the potential significance of the butyrate kinase pathway in the elevated levels of butyrate in these individuals (Fig. 4A). However, this increased elevation did not reach statistical significance.

Significantly, both butyrate kinase and acetate-CoA transferase play crucial roles in the production of butyrate. Notably, the abundance of butyrate kinase exhibited a statistically significant increase across all three age groups in individuals with a high methanogen phenotype (Supplementary Table). Upon closer examination, we observed a positive correlation between the genes responsible for butyrate production through the butyrate kinase pathway and members of Oscillospiraceae. In contrast, these genes showed negative correlations with members of Lachnospiraceae (Fig. 4B, Supplementary Fig. 8). This observation supports our hypothesis that individuals with a high methanogen phenotype, particularly as they age, may compensate for the reduction in butyrate levels by harboring elevated levels of Oscillospiraceae, particularly through the butyrate kinase pathway.

**Supplementary Fig. 8.**
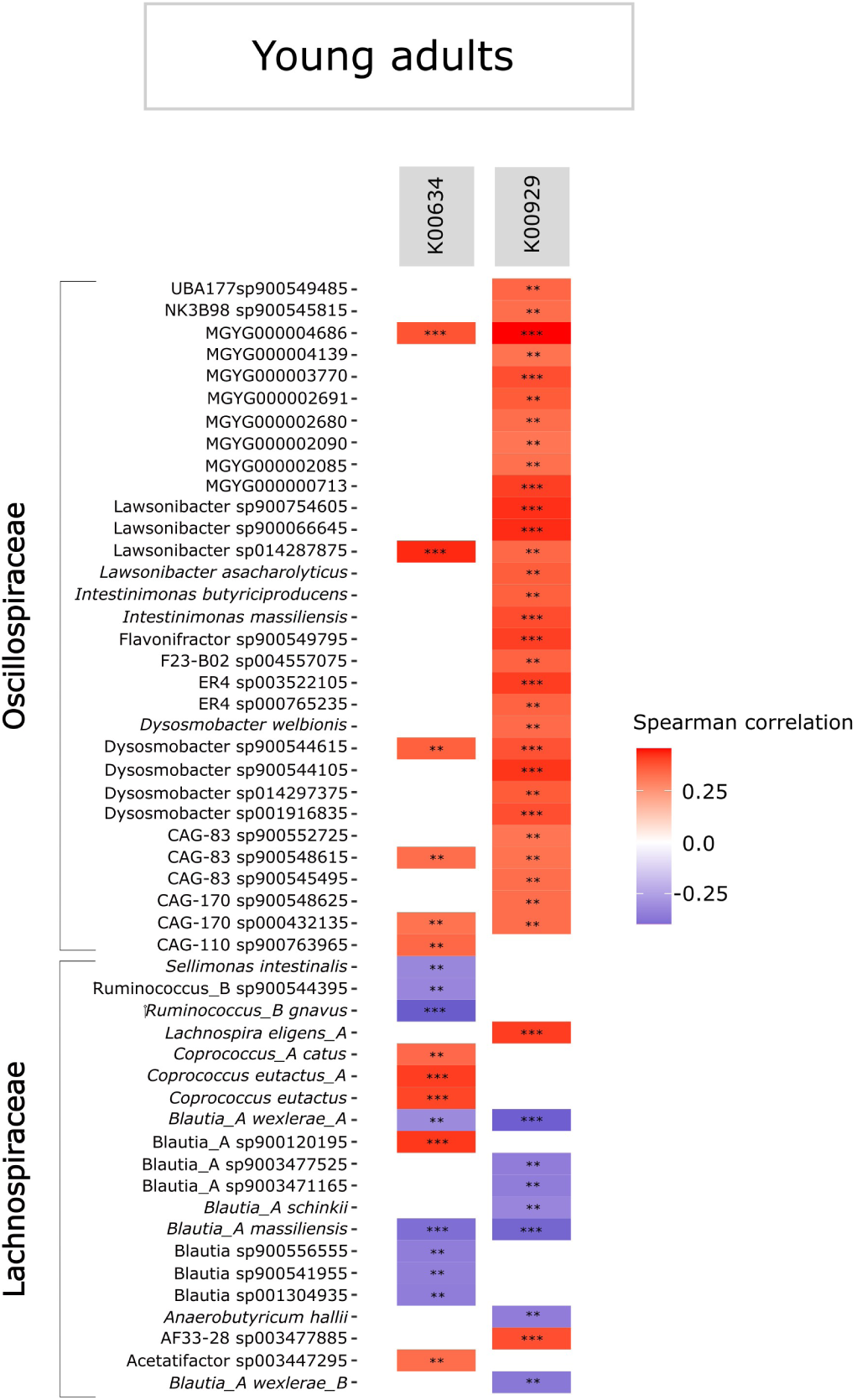
Correlation of butyrate kinase pathway genes in young adults with bacterial taxa. ***q<0.001, **q<0.01, *q<0.05.

## Discussion

Aging-associated changes in the gut microbiome have been tackled in several studies (61), however, despite the vital importance of the gut archaeome, there is a significant gap in our understanding regarding how age, and especially longevity, affects the distribution of archaea within the gut and *vice versa*. Specifically, the role of methanogenic archaea with a profound impact on the structure and functionality of the entire gastrointestinal microbiome remains poorly elucidated. Moreover, since *Methanobrevibacter smithii*, as the predominant archaeal species within the human gut, has been recently divided into two clades, namely *M. smithii* and *Ca.* M. intestini, highly resolved investigation of gut archaeome helps to deepen our understanding of the importance of these two archaeal species in longevity and their role in the emergence of high methanogen phenotype. Besides, how to thoroughly profile the composition of gut archaeome and define the high methanogen phenotype based on metagenomics data remains a challenge. The main goal of this study was to explore the gut archaeome of different age groups and to assess how the distribution of high methanogen phenotype changes with age.

Our study revealed that, with aging the gut archaeome richness and/or diversity decreased, which was in contrast to the previous reports based on 16S rRNA gene sequencing (15). We then delved more into the differences of the archaeal profile of different age groups. The increased relative abundance of *M. smithii* with advancement of age, and the possible increased abundance of this methanogen prior to longevity has been previously reported (62). Our study however provided evidence that although there is a steady increase in relative abundance of Methanobacteria with the progression of age, there is a transition phase where the OAs are characterized by the increased abundance of *Ca.* M. intestini compared to YAs and CENT rather than *M. smithii*. On the other hand, both YAs and CENT harbor a comparably high abundance of *M. smithii* in their gut. These results were in fact in contrast to the reported enrichment of *M. smithii* in CENT compared to YAs and OAs (47, 48), however, this discrepancy potentially arises from overlooking *Ca.* M. intestini as a separate archaeal clade and different methods of the profiling of archaea based on the genomic sequences.

Thermoplasmata is another commonly found archaeal taxon in the human gut that tends to be more abundant in OAs. This has led to the hypothesis that since these archaea have environmental origins, age could play a role in promoting their survival in certain individuals (16). Our observations also indicated the high abundance of these archaea in OAs compared with YAs, however, interestingly, the relative abundance of Thermoplasmata in CENT showed reduction as compared with OAs and was observed to be rather comparable to YAs. There is evidence suggesting that Thermoplasmata could potentially contain genes responsible for producing certain metabolites such as methylglyoxal, indole, and acetaldehyde with the potential of disrupting DNA or signaling pathways (63, 64). Although further molecular experiments are required to fortify the involvement of these archaea in disease progression, there might be a potential link between the presence of Thermoplasmata and disease, which could explain why they appear to be less abundant in individuals with a longer lifespan. Interestingly, according to the literature, within this archaeal order, certain species may counteract trimethylamine (TMA, involved in the progression of atherosclerosis), while some species lack this capability. On the other hand, the high abundance of *Ca*. Methanomassilicocales intestinalis in frail individuals raises questions about its role and it is not clear whether its high abundance is favored due to factors like altered gut transit or independently contributing to inflammation. Further research is needed to assess the immune potential of dominant Methanomassiliicoccales, especially those using TMA, and determine if their role is primarily positive through TMA removal or more nuanced (49).

According to the archaeal abundance in high methane emitters, with a significantly higher abundance of *Methanobrevibacter* (≥ 0.03%), subjects could be stratified into low and high methanogen phenotypes. Our study revealed that the prevalence of subjects with high methanogen phenotype increases with age. Interestingly, not only *M. smithii,* but also *Ca.* M. intestini was found to contribute to the emergence of high methanogen phenotype, with a more evident contribution of *Ca.* M. intestini in OAs, which can be argued by the increased abundance of this archaeal species in this age group. The exact cause of the link between the higher prevalence of high methanogen phenotype and the aging process remains somewhat elusive, however, the slower digestive transit times often observed in aging individuals, as well as their differences in dietary habits and contact with livestock could contribute to the overrepresentation of archaea (65, 66).

Subjects with high methanogen phenotype showed different microbiome signatures and higher microbial diversity, especially evident in YAs and OAs rather than CENT. Interestingly, a significantly higher alpha diversity has been frequently linked to improved stability and resistance to disruptions (67). This increased diversity of microbial species in subjects with high methanogen phenotype has been previously linked to the ability of methanogens to reduce the hydrogen partial pressure and thus facilitating the microbial fermentation, which is otherwise restricted by the hydrogen accumulation and inhibition of NAD coenzyme regeneration (68). In CENT with high methanogen phenotype, only the richness index was significantly higher and not the evenness. In general, despite occasional contradictions (69), lower gut microbial alpha diversity has been shown in CENT compared to that of YAs and OAs (12, 70). A variety of confounding factors can influence the controversial reports regarding microbiota alpha diversity with respect to aging. These factors include host and/or lifestyle factors as well as geography or the number of included subjects in cohorts. Moreover, although the CENT cohort represents a healthy population, the process of aging, particularly in its later stages, is associated with a natural decline in gastrointestinal function and the host immune response. This decline may contribute to the onset of chronic low-grade inflammation and metabolic disorders. (48, 71, 72). Therefore, a reduction in alpha diversity measures of high methanogen phenotype in CENT compared to those within YAs and OAs is not surprising.

When examining the relationship between the bacterial taxa associated with *M. smithii* or *Ca.* M. intestini in different age groups, we observed that in CENT, these species had more complex networks with other bacterial taxa compared to YAs and OAs, which was more akin to those seen in individuals with a high methanogen phenotype, who showed a dynamic archaea-bacteria network. Upon closer examination of these networks, we found that *M. smithii* or *Ca.* M. intestini consistently co-occurred regardless of the age group, which was in contrast to previous findings based on Sanger sequencing (73). Additionally, the family Christensenellaceae was consistently associated with these archaeal species across all age groups. The co-occurrence of *Christensenella* with *M. smithii* or *Ca.* M. intestini was consistent with previous research that demonstrated a mutually beneficial relationship between *Christensenella* and *Methanobrevibacter* through interspecies hydrogen transfer (74). *Methanobrevibacter* spp. play a crucial role in efficiently digesting complex polysaccharides by optimizing hydrogen levels for bacterial polysaccharide digestion and consuming the end products of bacterial fermentation. In our analysis of bacteria-archaea correlations, we identified Oscillospiraceae known for butyrate production (75, 76) and subsequent anti-inflammatory properties to co-occur with both *M. smithii* and *Ca.* M. intestini. This suggests a mutualistic or syntrophic relationship between these gut bacteria and archaea. Conversely, we observed a significant negative association between these archaeal species and members of the Lachnospiraceae family, which are also known butyrate producers in the gut (77). The mutual exclusion between Lachnospiraceae and methanogens is not surprising, given that Lachnospiraceae functions as acetogens, utilizing hydrogen and carbon dioxide in the gut to produce acetate. In contrast, methanogens employ these two substrates for methanogenesis. Consequently, these microorganisms engage in a competitive relationship for substrates (78, 79).

Interestingly, the cumulative presence of Lachnospiraceae, recognized as butyrate-producing bacteria, decreases with age (13). Our findings reveal that individuals with a high methanogen phenotype in the OAs and CENT age group exhibit elevated levels of genes (metagenome) associated with the butyrate production pathway, particularly the butyrate kinase pathway. Our correlation analysis highlights a positive association between the gene responsible for butyrate kinase, a pivotal enzyme in the butyrate production pathway, and members of Oscillospiraceae. This underscores the significance of these microbial taxa in butyrate production, a known health-promoting factor, among individuals with a high methanogen phenotype. Hence, it can be inferred that these individuals may compensate for the decline in Lachnospiraceae not only through the co-occurrence of *M. smithii* and Ca. *M. intestini* with Oscillospiraceae but also through an increased abundance of the latter in their microbiota. Consequently, it is plausible to suggest that, as individuals age, the reduction in Lachnospiraceae (potentially due to dietary factors) could be more manageable for those with methanogen phenotype, as indicated by comparable butyrate levels to the microbiome of subjects lacking this phenotype. Therefore, disruptions in the establishment of the high methanogen phenotype during aging could be a critical factor influencing the maintenance of optimal butyrate levels in the gut.

Another interesting observation was the consistent negative co-occurrence of *M. smithii* or *Ca.* M. intestini with *Streptoccocus*, especially in OAs. It is noteworthy to mention that increased levels of metabolic makers of dysregulation has been associated with increased abundance of *Streptococcus* and therefore it is mostly linked with unhealthy aging (80), suggesting that the presence of these methanogens could be linked to the advancement of OAs towards achieving healthy aging, rather than the progression of disease.

There are some shortcomings in our study. First, the food intake of individuals as well as their chewing ability with the progression of age could play an important role in the microbiome and archaeome compositions. However, we did not have access to the food questionnaire of the centenarians. Second, we did not have a longitudinal setting, where we could investigate at which age the enrichment of methanogens begins in the human gut. Finally, although we used a most recent database (including all genomic information from isolates and metagenome-assembled genomes for archaeal classification, there are still a large number of under-discovered and uncultivated nature of human colonized archaea. This could potentially result in an insufficient representation of archaeal taxa within the reference archaea database, consequently leading to incomplete profiling of fecal archaeal species diversity across different populations in the current study.

This study again supports the relevance of the archaeal microbiome component on human physiology and thus healthy aging. Our research highlights the dynamic, age-related shifts in methanogen composition, particularly the increasing prevalence of the high methanogen phenotype with advancing age. Our study emphasizes the significance of *Ca.* M. intestini, evident in its surge during the transition phase represented by older adults, its co-occurrence with *M. smithii*, and its substantial role in both methane emission and the emergence of the high methanogen phenotype. This is the first insight into a critical role of this new archaeal representative for the human host. Moreover, our findings underscore the importance of methanogens partnering with specific butyrate-producing bacteria using the butyrate kinase pathway, enhancing the health status of individuals and potentially contributing to longevity in those with a high methanogen phenotype.

## Acknowledgements

We express our gratitude for the computational resources provided by the MedBioNode at the Medical University of Graz, funded through the Austrian Federal Ministry of Education, Science, and Research, specifically under the Hochschulraum-Strukturmittel 2016 grant as part of BioTechMed Graz. Additionally, we appreciate the support extended by the ZMF team at the Core Facility Computational Bioanalytics, located at the Medical University of Graz. RM was supported by the local PhD program MolMed.

## Authors’ contributions

RM did the DNA extraction and data analysis, produced most of the figures and wrote the manuscript. AM supported data analysis and study outline. TS supported data analysis. VW contributed to figure preparations. CK and HS performed sampling. CME supervised all activities, performed analyses and wrote the manuscript. AM, TS, CK, and VW contributed to the writing of the manuscript.

